# Segmentation boosting with compensation methods in optical coherence tomography angiography images

**DOI:** 10.1101/2020.08.20.258905

**Authors:** Yih-Cherng Lee, Jian-Jiun Ding, Ling Yeung, Tay-Wey Lee, Chia-Jung Chang, Yu-Tze Lin, Ronald Y. Chang

## Abstract

Optical coherence tomography angiography is a noninvasive imaging modality to establish the diagnosis of retinal vascular diseases. However, angiography images are significantly interfered if patients jitter or blink. In this study, a novel retinal image analysis method to accurately detect blood vessels and compensate the effect of interference was proposed. We call this the patch U-Net compensation (PUC) system, which is based on the famous U-Net. Several techniques, including a better training mechanism, direction criteria, area criteria, gap criteria, and probability map criteria, have been proposed to improve its accuracy. Simulations show that the proposed PUC achieves much better performance than state-of-art methods.

## Introduction

Retinal vasculature analysis is a critical subject in diagnosing and managing retinal vascular diseases. Optical coherence tomography angiography (OCTA) is an advanced rapid noninvasive method that can acquire the images for retinal vasculature analysis [1–4]. Chorioretinal vasculature contains a three-dimensional layered structure and it comprises a superficial layer, a deep layer, an outer retinal layer, and a choriocapillary layer. OCTA facilitates the detection of retinopathy [5], glaucoma [6], and diabetic complications [7]. However, OCTA images can be affected by human interference. For example, patients may blink unconsciously during OCTA infrared scanning. Therefore, it is common for an OCTA image to product several megascopic artifacts. Additionally, the vasculature sometimes tends to be rare, small, and thin due to some disease. These problems may degrade the quality of an OCTA image. In this study, we tried to solve the following two problems for OCTA image analysis:

i. The image quality problem of noise-interfered blood vessels: The noise may be caused by the environment, equipment, the vessel in the next layer, and small nerve fibers in the plexus. Additionally, as indicated previously, the motion of a patient may produce flicker noise.
ii. Disease problem: In a healthy retinal image, vessels are evenly distributed on the retina. However, in a diseased retinal image, severe vascular non-perfusion may occur. There are large empty / black areas, making it difficult for a doctor to identify whether a cluster of bright pixels in the dark region is indeed a vessel or an artifact.

Considering the above problems, some advanced noise filters and thresholding methods have been proposed. For example, the adaptive thresholding methods proposed in Phansalkar et al. [8] and Cole et al. [9] and the multiple enface image averaging Uji et al. [10] were well-known methods for image intensity enhancement. Moreover, to reduce the noise and enhance the visual ability of the retina, in [11–18], several techniques were adopted, including the Gabor-filter-based method [11], the Frangi-filter-based methods [12], the filter-bank-based architecture [13, 17], the optically-oriented-flux [14] and thresholding [18]. In [15], the compressive sensing method was applied to remove the noise of OCTA images. In [16], the generalized Gauss-Markov random field and the guided bidirectional graph search method were applied to perform retinal vessel segmentation in OCTA images, respectively. However, these methods are not unsupervised and parameter-sensitive and may produce unexpected noise by personal setting operations. In recent years, with the fast development of deep learning, several learning-based segmentation methods, including the convolutional neural network (CNN)-based [19] and the U-Net-based methods [20, 21] have been proposed. These methods can reproduce the curvilinear vessel shape for healthy retinas. However, when handling diseased retinal images, there are some limitations and the performance is affected by the artifact and the non-uniform distribution of vessels. Therefore, in this study, we integrated the advantages of deep learning networks and conventional methods and proposed the patch U-Net compensation (PUC) algorithm. The main concepts of the proposed PUC algorithm are summarized as follows:

a. To solve the vessel segmentation problem: We modified the original U-Net model [20] by varying its training process to make the model fit OCTA images perfectly. This modified U-Net model is treated as the backbone of the proposed PUC algorithm. It can well identify the vessels with large curvature.
b. To solve the diseases retinal image problem: We developed several techniques for artifact identification and vessel compensation to address the problem well.

First, we proposed an artifact identification technique based on corner detection, edge detection, and ridge detection [22]. A novel direction estimation method and area thresholding method were adopted to identify whether a cluster of bright pixels was a real noise or a small vessel. Its idea is based on human vision. It applies the fact that the vessel should be ridge-like and fiber-like to distinguish the artifacts and small vessels.

Subsequently, an adaptive compensation technique was applied to the remaining vessels. The U-Net [20, 21] performs vessel identification globally since its loss function is determined from whole pixels, not for the pixels in a special region. Therefore, we proposed a region-adaptive technique to connect small fragments in suspected regions to refine the output locally. These compensation techniques will be illustrated in detail in Sections “Compensation methods for noise reduction” and “Vessel compensation”.

## OCTA images and challenges in analysis

RTVue XR 100 Avanti Edition [23], which is an OCTA machine, provides the scanned images from the retina by the reflections on different vessels, as shown in Fig 1(f), to obtain four-layer information, as shown in Figs. 1(a)(b)(c)(d). Retina vessels are distributed like tree branches. Larger vessels (i.e. arterioles and venules) are mainly found in the superficial layer, while small vessels (capillaries) can be found in both superficial and deep layers. Vascular non-perfusion in superficial layer is an important feature of diagnosing and evaluating retinal vascular diseases.

**Fig 1.**
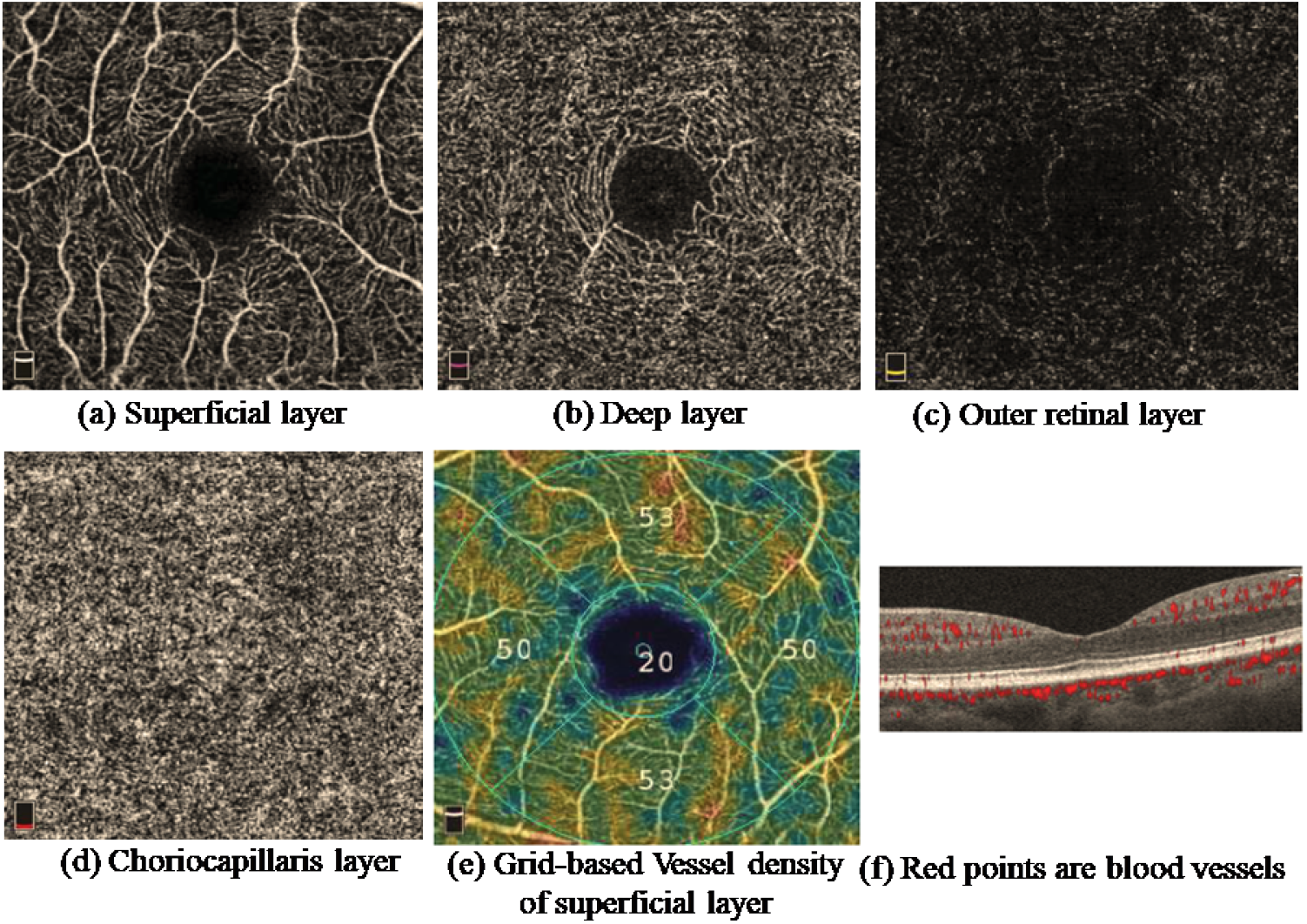
Outputs of the OCTA machine of RTVue XR 100 Avanti Edition, including (a)(b)(c)(d) four layers, (e) the vessel density map, and (f) cross-sectional images. The red points indicate vessels.

Although OCTA machine may provide a vessel density map [24–27], as shown in Fig 1(e), the result is not robust to artifacts and noise, which may be misidentified as small vessels. How to filter the noise, like the red arrows, as shown in Fig 2(a), and retrieve the true vessels, as shown in Fig 2(b), is a challenging problem. Even when the patient moves the head slightly or blinks a little, noise is produced. Moreover, the scanning artifact problem may form a straight line in an OCTA image. It is often misidentified as a vessel. Consequently, it is insufficient to use only the intensity information to distinguish between vessels and noise. In particular, if the intensities of scanning artifacts are high and the vessels are blurred, using only the intensity information to identify vessels may cause several errors. Therefore, instead of applying some simple rules for vessel detection, a more sophisticated method is required.

**Fig 2.**
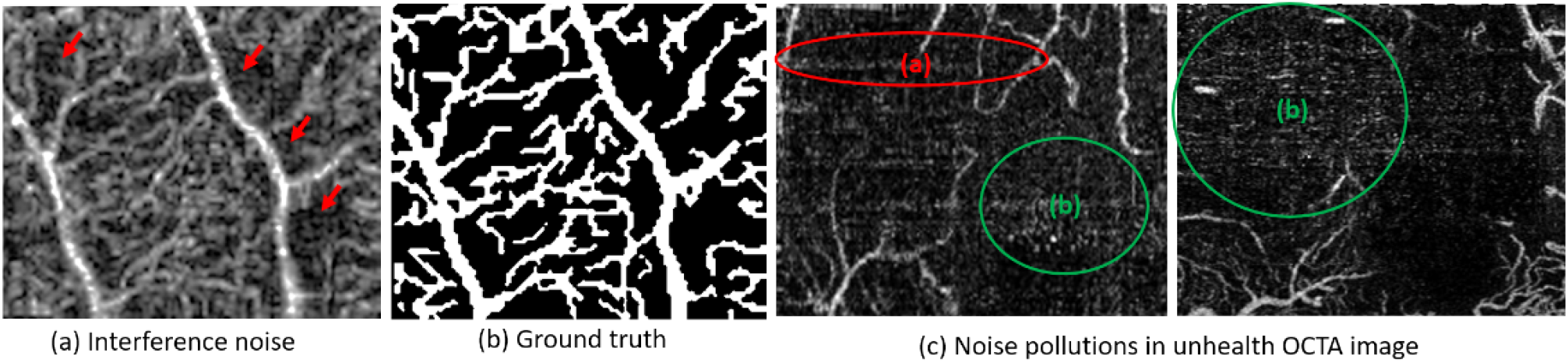
(a) An OCTA raw image with noise (marked by red arrows); (b) the ground truth; (c) the scanning artifact problem (red ellipse) and noise (green ellipses) may be misidentified as vessels and affect the performance.

In an OCTA image, if there is some bright region that is in fact not a vessel, then it is treated as noise. Due to machine oscillation and the motions of patients, most noise is either pepper-like or white-noise-like. Because the vessels, which are usually ridge-like, have several high-frequency components, it is suitable to apply the frequency-based filter (e.g., the lowpass filter) to remove the noise. Several studies [8–21] have been proposed to remove the bulk motion noise problems, as shown in Fig 2(a). However, it is still a challenge to deal with the strong artifacts and straight artifact lines, as shown in Fig 2(c).

In Fig 2(c), the red circles represent the strange thin straight line which comes from blink interference during the horizontal infrared scanning procedure and the green circle represents the noise interference from the OCTA machine. In this study, we used the direction criteria and the area criterion in Section “Compensation methods for noise reduction” to remove this type of noise.

## Materials and Methods

The U-Net, which is a multilayer deep learning model, has been applied in medical image processing [20, 21]. For example, in [20], the U-Net was applied to neuronal membrane image processing. However, OCTA images have denser vessel distributions and the noise interference problem is more severe in OCTA images than in other medical images. Therefore, in this study, we proposed an advanced method based on modifying the U-Net model to achieve a better vessel extraction result. The architecture of the proposed algorithm is plotted in Fig 3. It comprises two parts: (i) the patch U-Net and (ii) compensation methods.

**Fig 3.**
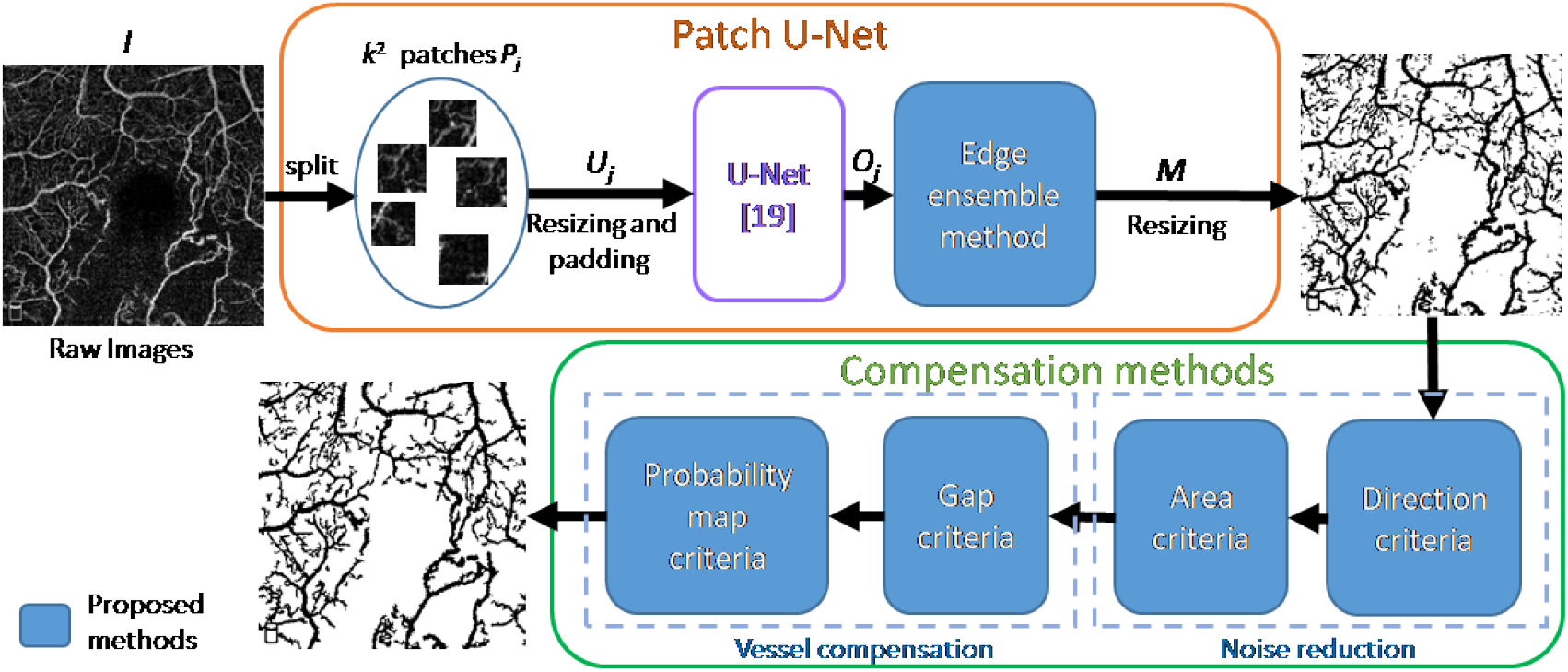
Architecture of the proposed PUC system.

### Patch U-Net

The U-Net is widely used for segmentation. However, when we adopted it directly on OCTA images, the performance may be limited. If one feeds the original OCTA image into the U-Net directly, the training process may not be convergent to a good vessel detection result. Furthermore, taking the whole OCTA image may increase the use of the GPU memory. Because OCTA images are complicated and include several tiny vessels and artifact interferences, we found that using the U-Net directly could not describe the details of tiny vessels. Most of time, the model often ignores both tiny vessels and artifacts.

To deal with this problem, an interesting patch-based training strategy was proposed in this study. We also developed a meaningful and significant patch training strategy in OCTA image training on the U-Net-based model. We divided the input OCTA image into small and fixed size patches. Considering that the vessels in OCTA images were relatively small, using the proposed patch-based training strategy is very helpful for detecting tiny vessels successfully without increasing the effect of noise.

After obtained the outputs of the U-net for all patches, we fused these results. However, if one performs fusion directly, the discontinuous edge problem may occur at the boundaries of patches, as shown in Fig 4(b). Therefore, we proposed a patch ensemble method to address this issue. We applied the following methods to ensure that the output had continuous edges.

**Fig 4.**
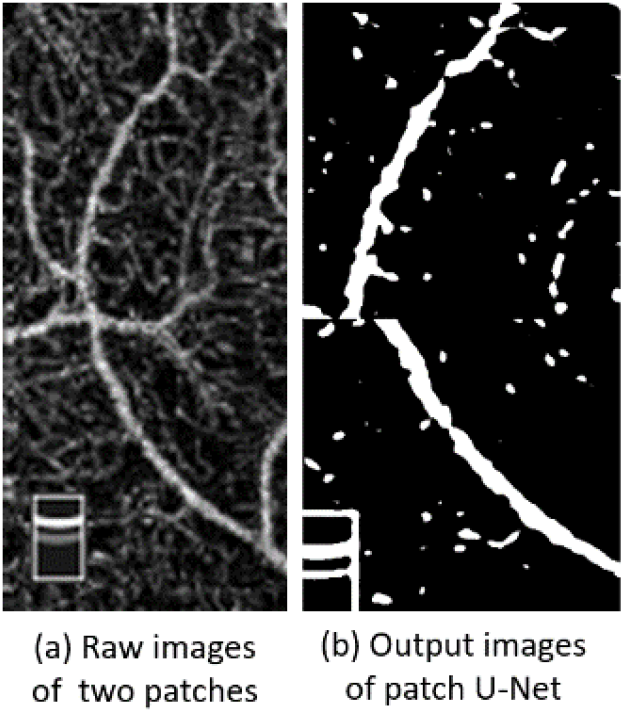
Illustration of the problem in the assembling process of the patch U-Net.

First, suppose that *H* is the input size of the U-Net, *Ĥ* is the output size of the U-Net, and *k*×*k* is the number of patches divided from the input image *I*. We padded each patch before network prediction to avoid the edge truncation effect. According to Algorithm 1, we generated, padded, and resized patches and used them as the inputs for U-Net training.

Second, we input a set of new patches *U_j_*(*j* = 1, 2,…, *k*×*k*) to the U-Net model to acquire accurate output *O_j_*(*j* = 1, 2,…, *k*×*k*) for the final ensemble procedure, as shown in Algorithm 2. Then, we resized the outputs and combined these by orders of Algrithom1 division orders.

The hyper-parameters are as follows: the input OCTA image size is *N* = 304, *H* is 572, *Ĥ* is 388, and *k* = 4. We divided the raw image into 16 uniform patches with size of 76×76. Then, each patch is symmetrically padded and treated as the input of the U-Net.

**Algorithm 1:**
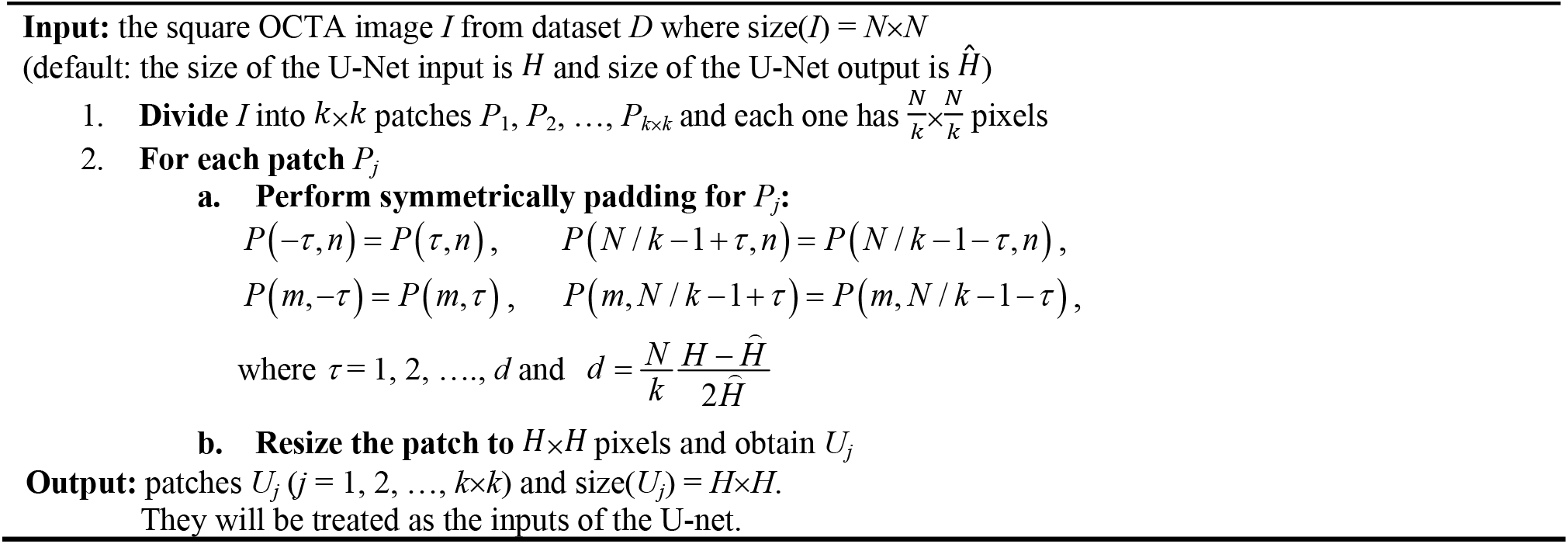
Generate Patches

**Algorithm 2:**
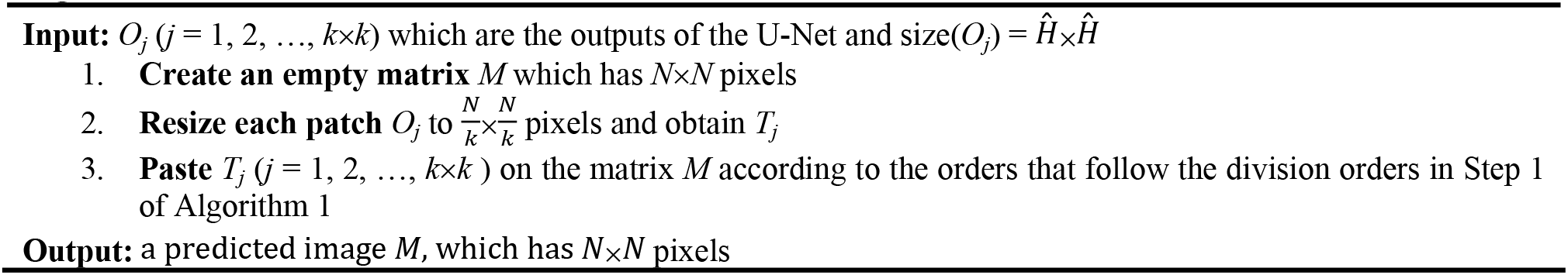
Patch Ensemble Methods

### Compensation methods for noise reduction

After applying the patch U-Net to determine vessels from the retinal image, the output was compensated to distinguish between the noise and true vessels. The proposed compensation method comprises two parts: (a) the noise reduction part and (b) the vessel compensation part. The noise reduction part applies two methods, (i) direction criteria and (ii) area criteria, to refine the output of the patch U-Net model and remove the noise and artifact parts. The detail of the vessel compensation part will be described in Section “Vessel compensation”.

i. Direction criteria: First, the output of the patch U-Net may contain some fragments and noise, as shown in Fig 5(a). We removed them according to their unique features. We can apply the techniques of feature point classification, which can classify a point into a peak, a ridge pixel, an edge pixel, a corner, or a flat pixel [22]. For each pixel, we observed the variations along the eight local directions, which were denoted by **e1**, −**e1**, **e2**, −**e2**, **e3**, −**e3**, **e4**, and −**e4**. To determine **e1**, first, we computed the convolution with the kernel matrix **K** and took the angle:

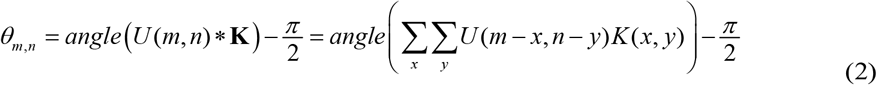

where * means convolution, *U*(*m, n*) is the output of the patch U-Net, and

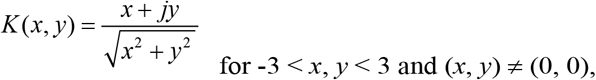 *K*(*x, y*) = 0 otherwise. Then, **e1** was determined from

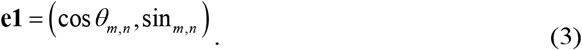 Note that **e1** varies with the location (*m, n*). Second, we performed clockwise rotation for **e1** with 45°, 90°, and 135° to produce **e2**, **e3**, and **e4**, respectively, as shown in Fig 5(c). Then, we computed the variations along the directions of ±**e1**, ±**e2**, ±**e3**, and ±**e4**, respectively. For example, to compute the variation along **e1** for pixel (*x*, *y*), we first set

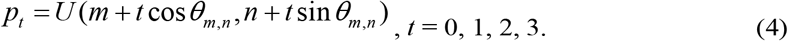 If *m*+*t*cos*θ_m,n_* or *n*+*t*sin*θ_m,n_* is not an integer, then bilinear interpolation will be applied. Then, the variation along **e1** (denoted by *V*_1_) was determined from the weighted sum of *p_n_*–*p*_0_:

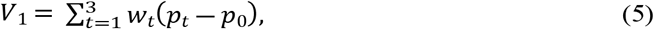

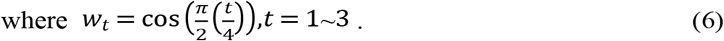 The variations along **e2**, **e3**, **e4**, −**e1**, −**e2**, −**e3**, and −**e4** (denoted by *V*_2_, *V*_3_, *V*_4_, *V*_-1_, *V*_-2_, *V*_-3_, and *V*_-4_, respectively) can also be determined from Eq (4)–Eq (6) but *θ_m,n_* in (4) is replaced by *θ_m,n_* + *kπ*/4 where *k* = 1, 2, 3, −4, −3, −2, and −1, respectively.
ii. Area criteria After denoising by the direction criteria, we tried to remove the artifacts. Different from noise, the artifact is not an isolated dot and cannot be removed by the direction criteria. However, we found that the artifact usually has a very small area. For a 304×304 image, if a region with an area smaller than 7 pixels, it could be absolutely defined as an artifact. Therefore, we use a well-known contour algorithm [28] to determine all isolated regions and filter the regions whose area is less than 7 pixels.

**Fig 5.**
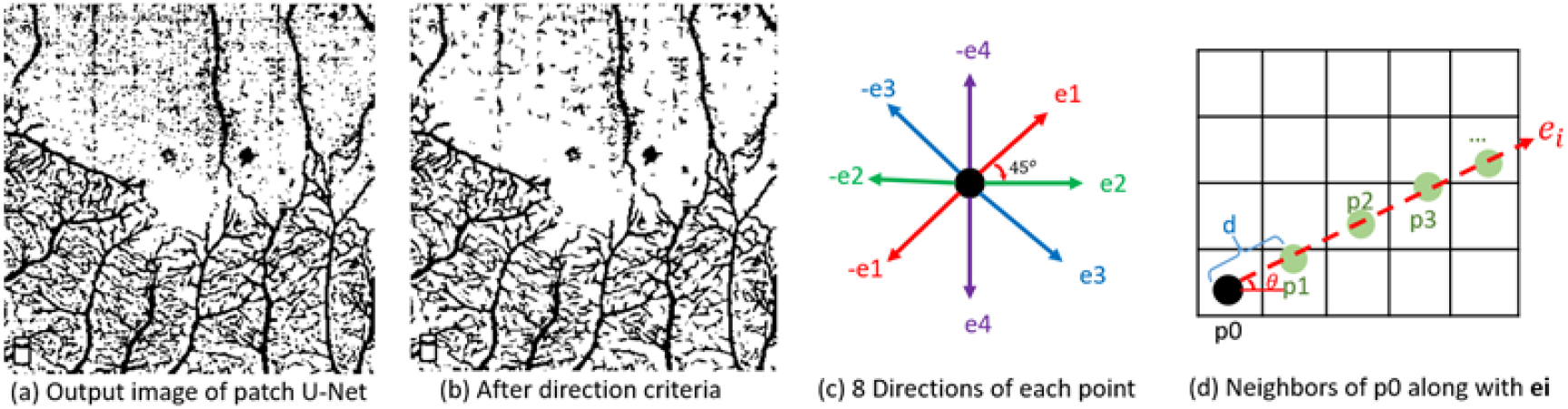
Illustration of the method of direction criteria.

### Vessel compensation

In addition to noise and artifact removal, we also propose several techniques for fragment connection. In a blurred OCTA image, sometimes some parts of the vessel are not well detected and a vessel will be broken into several fragments in the detection output. Because a vessel should have a line shape, after performing the patch U-Net, it is appropriate to connect the two fragments that have a small gap, as shown in Figs. 6(b)(c). Therefore, we proposed a geometric method to connect vessel fragments.

iii) Gap criteria If a group of segments in the detection result corresponds to the same vessel in the ground truth, then they have line shapes and their directions and locations should be similar to those of some surrounding fragments. By contrast, noise and artifact parts are usually isolated dots. Although the noise from the scanning artifact also has a line shape, it is very thin and too straight and can be well removed by the U-Net. Therefore, we used the famous principal component analysis method to determine the principal axis of the smaller fragment, as shown in Fig 6(d). The normalized principle axis is denoted by **e1** and the direction orthogonal to **e1** is denoted by **e3**. Then, we determined the line that could connect the end points of the two fragments. If the projections of the line on **e1** and **e3** are *α* and *β*, respectively, and

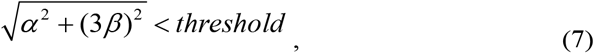

then the gap can possibly be a part of a vessel. In simulations, the threshold is chosen as 8, which is 1/38 of the image width. We used 3*β* instead of *β* to ensure that the connecting line had less projection on **e3**.
iv) Probability map criteria However, if one connects two fragments according to only the length of the gap, it is not enough to predict a more complicated association between two isolated fragments. Therefore, we also adopt the possibility map acquired from the output of the patch U-Net to determine whether two fragments should be connected. The original U-Net architecture always uses a fixed threshold (*th* = 0.5) to conclude whether a pixel belongs to the vessel or the background. However, it does not consider the association with neighboring regions. In the proposed algorithm, we adopted an adaptive threshold. For the gap between two fragments that satisfies Eq (7), more attention should be paid and we lowered the threshold for the pixels within the gap to make them easier to be identified as vessel pixels. Moreover, for the non-gap part, the possibility map of the U-Net is also beneficial for refining the vessel detection output. If *U*(*m, n*) and *A*(*m, n*) are the possibility map of the U-Net and the intensity of the original OCTA image, respectively, when *U*(*m, n*) + *A*(*m, n*)/800 ≥ *th_up_* = 104/160 then the pixel (*m*, *n*) is identified as a vessel pixel. When *U*(*m*, *n*) + *A*(*m*, *n*)/800 ≤ *th_down_* = 95/160, (*m*, *n*) is identified as a background pixel. If both the two inequalities are not satisfied, the criterion of *U*(*m, n*) > 0.5 is applied to determine whether a pixel is a vessel pixel.

**Fig 6.**
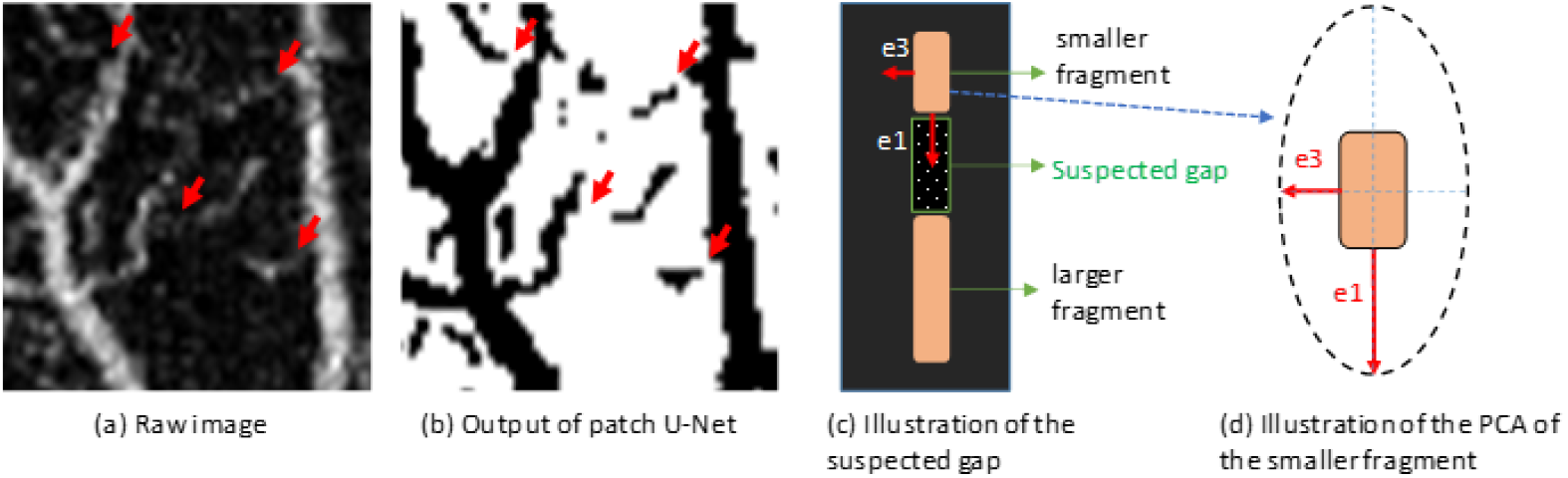
Illustrations of the gap criteria.

## Results

### Database

Our database was obtained from the Chang Gung Memorial Hospital, Keelung, Taiwan. The study followed the tenets of the Declaration of Helsinki and it was approved by the Chang Gung Memorial Hospital Institutional Review Board. All OCTA images used in this study were the superficial layer of retinal vasculatures and had a size of 304×304. Half of the images were from eyes with retinopathy and others were from healthy cases. As a practical clinical scenario, most images in the database were interfered with by noise or artifacts. It comprises 42 raw OCTA images with 42 annotated ground truth, including 17 training images, 17 validation images, and 8 test images. For the training network, we also applied data augmentations (flipping horizontally or vertically and rotating in degrees [-45, 45]).

### Evaluation

We use four metrics for evaluation: (i) accuracy, (ii) precision, (iii) recall (sensitivity), and (iv) the F1 score (Dice similarity coefficient). They are computed from the number of true-positive, true-negative, false-positive, and false-negative cases (denoted by TP, TN, FP, and FN, respectively):

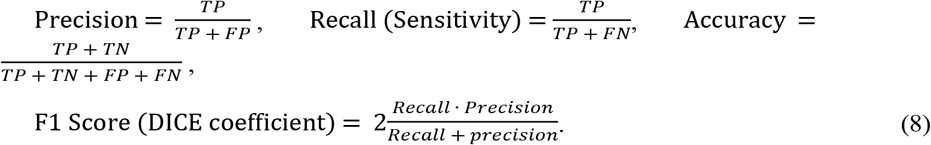

Among these metrics, the F1 score takes both precision and recall into consideration and can well reflect the performance of an algorithm.

### Experiments

In Table 1, we compare the proposed algorithm with several popular retinal vessel detection algorithms [11–14, 19, 21], including signal-processing-based methods [11, 12], traditional machine-learning-based methods [13, 14], and deep-learning-based methods [19, 21] associated with OCTA images. The results in Table 1 show that the proposed algorithm has a significantly better performance than the state-of-the-art methods. Compared to the study in [21], which is called the automated and network structure preserving segmentation (ANSPS) method, the F1 score of the proposed algorithm is 3.60% higher. Compared to other methods, the F1 score of the proposed algorithm is 5.17%-13.91 % higher.

**Table 1.**
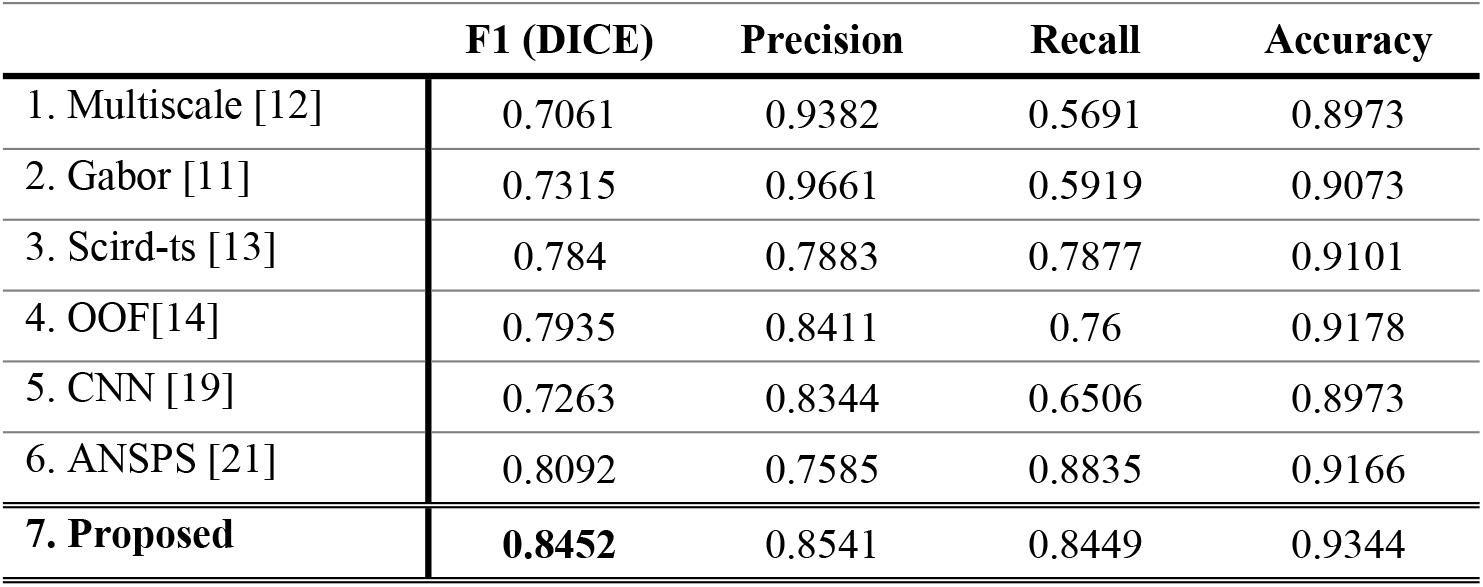
Performance Comparison for Vessel Detection in Retinal OCTA Images

|  | **F1 (DICE)** | **Precision** | **Recall** | **Accuracy** |
| --- | --- | --- | --- | --- |
| 1. Multiscale [12] | 0.7061 | 0.9382 | 0.5691 | 0.8973 |
| 2. Gabor [11] | 0.7315 | 0.9661 | 0.5919 | 0.9073 |
| 3. Scird-ts [13] | 0.784 | 0.7883 | 0.7877 | 0.9101 |
| 4. OOF[14] | 0.7935 | 0.8411 | 0.76 | 0.9178 |
| 5. CNN [19] | 0.7263 | 0.8344 | 0.6506 | 0.8973 |
| 6. ANSPS [21] | 0.8092 | 0.7585 | 0.8835 | 0.9166 |
| **7. Proposed** | **0.8452** | 0.8541 | 0.8449 | 0.9344 |

Moreover, in Table 2, we conduct an ablation study. We test the performance of the proposed algorithm where (i) only the patch U-Net is adopted, (ii) direction criteria are adopted, (iii) direction + area criteria are adopted, (iv) direction + area + gap criteria are adopted, and (v) all the proposed techniques (the patch U-net and the direction + area + gap + probability map criteria) are applied. The results in Table 2 show that the proposed direction and area criteria are indeed beneficial in improving the performance of vessel detection, since they can remove noise and artifacts. However, if one applies the gap criteria alone without using the probability map criteria, the F1 score is decreased. If both the gap and the probability map criteria are applied, the F1 score can be significantly improved.

**Table 2.** Ablation Study of the Proposed Algorithms

|  | **F1 (DICE)** | **Precision** | **Recall** | **Accuracy** |
| --- | --- | --- | --- | --- |
| (i) PUN | 0.7899 | 0.7128 | 0.9032 | 0.9028 |
| (ii) PUN+DC | 0.8315 | 0.793 | 0.8857 | 0.9244 |
| (iii) PUN+DC<br>+AC | 0.8376 | 0.8258 | 0.8594 | 0.9286 |
| (iv) PUN+DC<br>+AC+GC | 0.8313 | 0.8096 | 0.8648 | 0.9253 |
| **(v) PUN + DC**<br>**+ AC + GC + PMC**<br>**(Our PUC system)** | **0.8452** | 0.8541 | 0.8449 | 0.9344 |

<sup>1</sup>PUN, patch U-Net, DC, direction criteria, AC, area criteria, GC, gap criteria, PMC, probability map criteria, PUC, patch U-Net compensation

Moreover, in Figs. 7 and 8, we perform a visual comparison of the proposed algorithm and other vessel detection algorithms. Compared to traditional signal processing methods and machine learning methods [11–14], the proposed algorithm can well retrieve the details of vessels. Even if the widths of the vessels are very small, they can be retrieved well by the proposed algorithm. Compared to the CNN-based method, when using the proposed algorithm, the small vessels can be well detected and the widths of the detected vessels match those in the ground truth. Compared to the ANSPS method [21], the proposed algorithm is more robust to the interference of noise and artifacts. Moreover, the connectivity of the detected vessels of the proposed algorithm is better than those of other algorithms.

**Fig 7.**
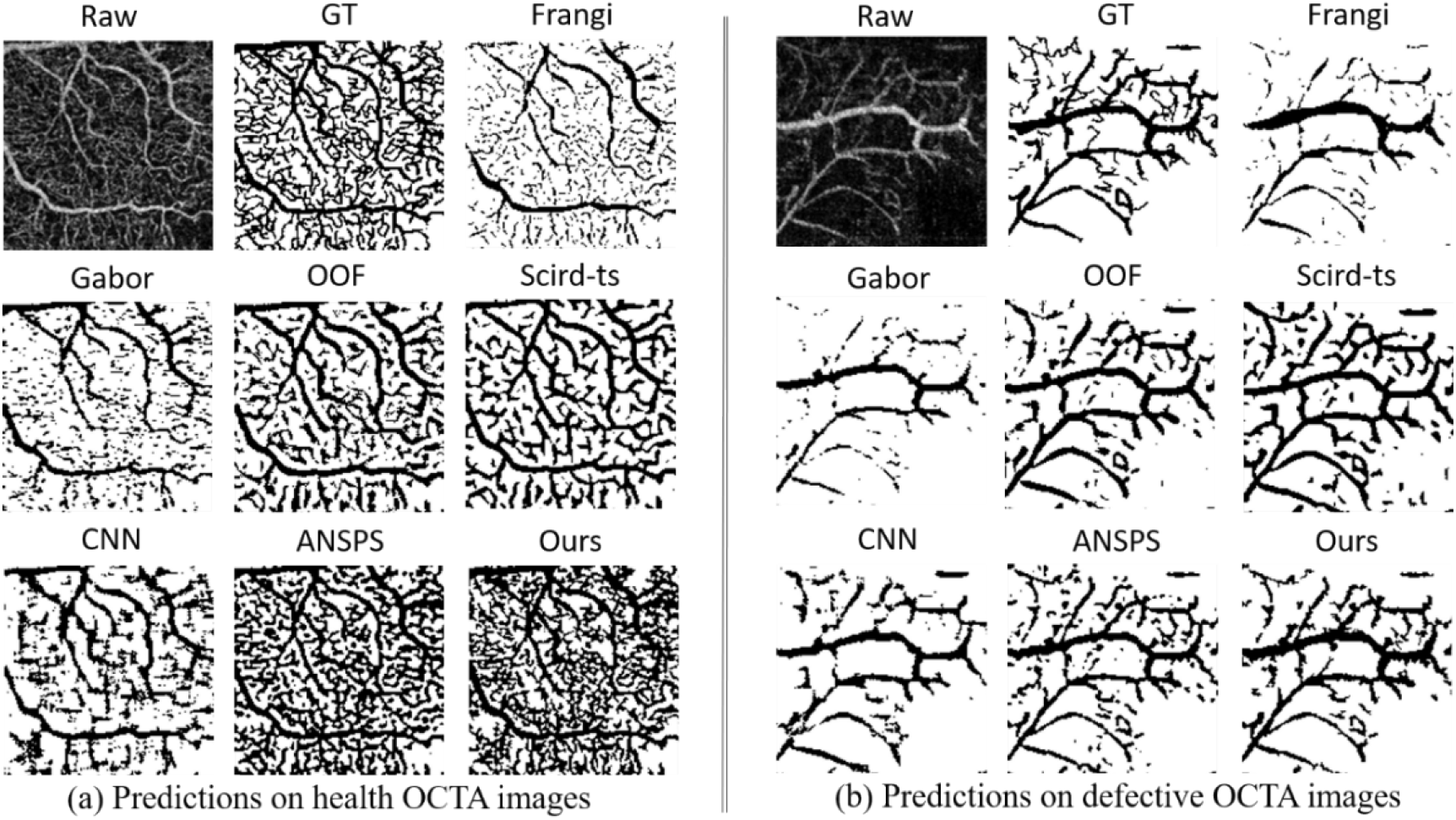
Illustration of the method of direction criteria.

**Fig 8.**
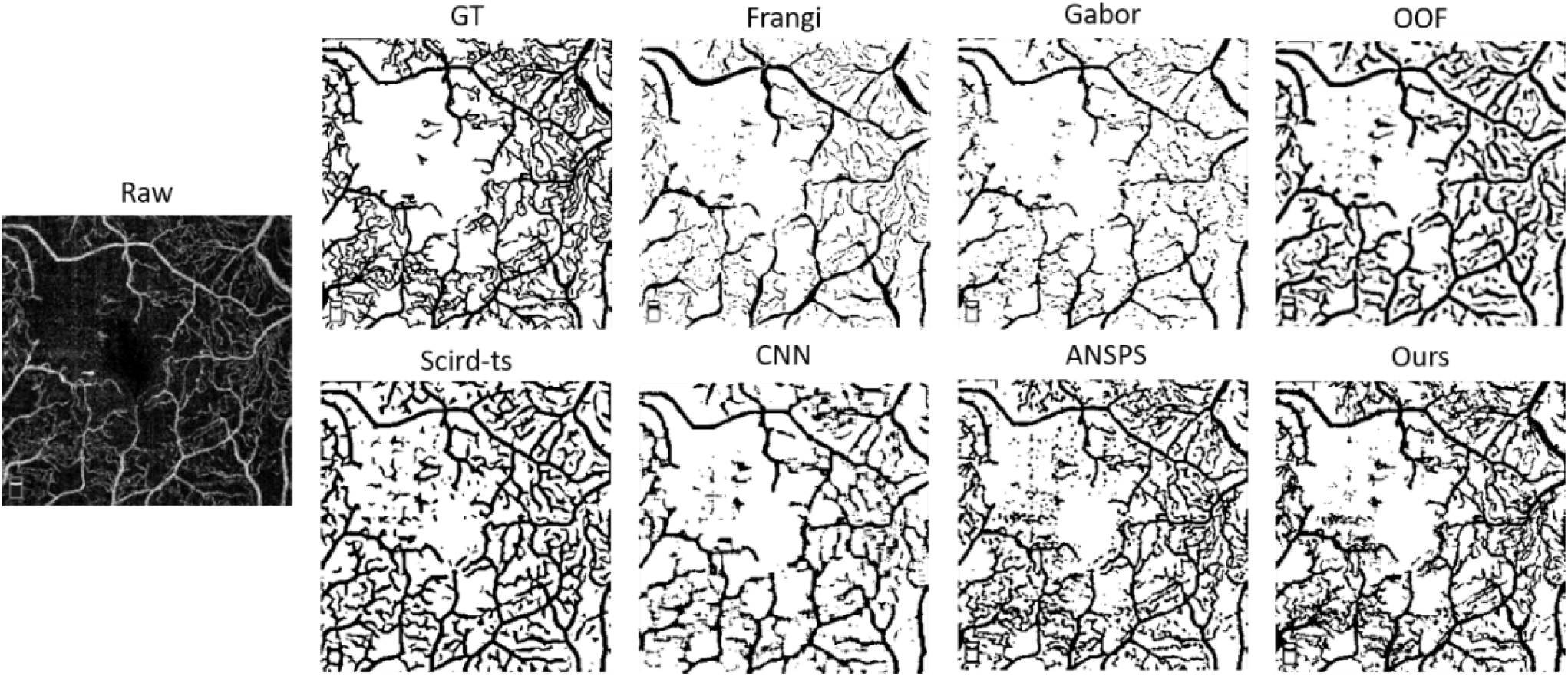
Illustrations of the gap criteria.

In retinal OCTA image analysis, it is important to retrieve the details of vessels while avoiding noise interference. However, there is usually a tradeoff between the two goals. With the proposed algorithm, both the two goals can be well achieved and high-quality vessel detection results can be acquired.

## Conclusion

A novel PUC algorithm has been proposed for blood vessel detection on defective OCTA images. It adopts various techniques, including the deep learning architecture, the noise and artifact removal mechanism using the direction and area criteria, and the vessel compensation mechanism using the gap and probability map criteria. The proposed algorithm effectively compensates the disadvantages of learning-based methods and can well identify whether a small bright region is a true vessel or artifact. Through the proposed compensation method, we do not require a large amount of training data as regular CNN-based methods. The proposed algorithm can facilitate the evaluation of the vessel density and is beneficial for retinal diagnosis.

## Acknowledgments

The authors thank for Keelung Chang Gung Memorial Hospital, Keelung, Taiwan, who helped us for data collection.

